# SciMind: A Multimodal Mixture-of-Experts Model for Advancing Pharmaceutical Sciences

**DOI:** 10.1101/2024.07.16.603812

**Authors:** Zhaoping Xiong, Xintao Fang, Haotian Chu, Xiaozhe Wan, Liwei Liu, Yameng Li, Wenkai Xiang, Mingyue Zheng

## Abstract

Large language models (LLMs) have made substantial strides, but their use in reliably tackling issues within specialized domains, particularly in interdisciplinary areas like pharmaceutical sciences, is hindered by data heterogeneity, knowledge complexity, unique objectives, and a spectrum of constraint conditions. In this area, diverse modalities such as nucleic acids, proteins, molecular structures, and natural language are often involved. We designed a specialized token set and introduced a new Mixture-of-Experts (MoEs) pre-training and fine-tuning strategy to unify these modalities in one model. With this strategy, we’ve created a multi-modal mixture-of-experts foundational model for pharmaceutical sciences, named SciMind. This model has undergone extensive pre-training on publicly accessible datasets including nucleic acid sequences, protein sequences, molecular structure strings, and biomedical texts, and delivers good performance on biomedical text comprehension, promoter prediction, protein function prediction, molecular description, and molecular generation.

## 1 Introduction

Large language models (LLMs) have made substantial strides, providing a versatile, task-agnostic base for a variety of applications[1], [2], [3]. However, their use in reliably tackling issues within specialized domains, particularly in interdisciplinary areas like pharmaceutical sciences, is hindered by several obstacles. These include data heterogeneity, knowledge complexity, unique objectives, and a spectrum of constraint conditions, which block the creation of groundbreaking applications[4], [5], [6]. This research aims to lay the groundwork for a large-scale model within the pharmaceutical sciences. In this area, four diverse modalities including nucleic acids, proteins, molecular structures, and natural language are involved. Of them, nucleic acids, proteins and molecular structures are the common modalities modeled by the pharmaceutical science community. Predicting the properties of a molecule or a protein[7], [8], [9], designing and optimizing for new ones[10], [11], [12], and understanding how they interact with each other [13], [14], [15] are common tasks and have made great progress. For example, AlphaFold3 and RosettaFold All-Atom models can even predict all interactions among these modalities.

However, a gap exists between these interactions and biological functions. While binding is common between proteins and molecules, the effects it may cause are rare and often expressed in natural language after experimentation, making standardization for modeling challenging. The effects a molecule can cause by binding to a protein are diverse, including competitive inhibition, non-competitive inhibition, agonizing, antagonizing, allosteric regulation, covalent modification, transport, and chelation, among others[16]. These effects are interconnected yet distinct from one another. Modeling each effect separately requires standardization and a separate classification or regression model, often leading to a loss of semantic meaning in the labels. In contrast, natural language descriptions provide an abstract and meaningful form of labeling for data, capable of conveying rich information.

Recent advancements in LLMs have propelled the development of cross-modal models between language and other modalities[4], [17], [17], [18], [19], [20]. These models, which include language-molecule, language-protein, and language-nucleic acids modalities, extend our capabilities to predict molecule functions, generate or optimize molecules with flexible constraints, annotate protein functions, and create or optimize proteins. However, their modality fusion is limited to two modalities.

In the field of pharmaceutical sciences, multiple modalities can be integrated, as depicted in Figure 1. If a model capable of managing all these modalities exists, then all biomedical text knowledge could be stored in a richly informative format. To address this, we’ve developed a specialized token set designed to individually tokenize different modalities. We also introduce a novel pre-training and fine-tuning strategy that harnesses the benefits of large-parameter models while minimizing their costs. This strategy, based on previous work MoEfication[21], involves two key components: (1) splitting the parameters of Feed-Forward Networks (FFNs) into multiple functional partitions called experts, and (2) building expert routers to determine which experts will be used for each input. By adopting a selective unequal number of expert activation strategy on different tokens, this approach enables data from different modalities to choose the most appropriate processing path. This approach not only results in a sparser model architecture, thereby reducing inference costs, but also circumvents modal alignment and potential performance decreases due to model size reduction. The main contributions of our work are as follows:

**Figure 1:**
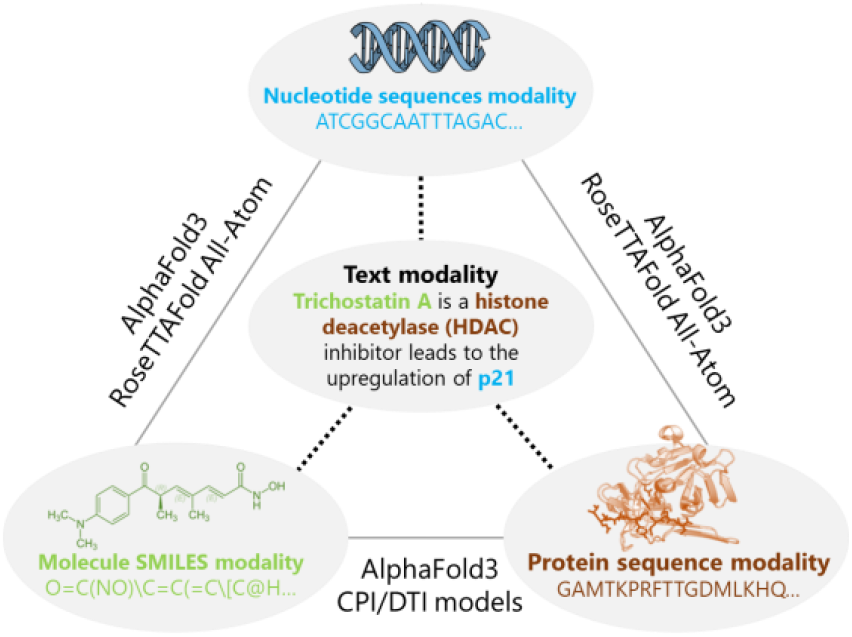
An overview of the four modalities in pharmaceutical sciences. The three traditional modalities, including nucleic acids (DNA/RNA), proteins, and small molecules, are typically modeled independently. Recent advancements have been made in the realm of cross-modal modeling, as indicated by the solid lines. However, there is a gap domain. In recent times, the natural language modality has surfaced as a highly promising method to describe nucleotide sequences, small molecules, and proteins, and it is swiftly garnering attention.

1. We’ve created a multi-modal mixture-of-expert foundational large model for pharmaceutical sciences, named SciMind. This model has undergone extensive pre-training on publicly accessible datasets including nucleic acids, protein sequences, molecular structures strings, and biomedical texts, and could be fine-tuned for downstream tasks involving all modalities in pharmaceutical sciences.
2. SciMind achieves state-of-the-art performance on benchmarks of molecular captioning and molecular generation by description.

## 2 Related works

In this section, we will provide a concise overview of the related work on cross-modal models in the field of pharmaceutical sciences.

### 2.1 Cross Language-Molecule Modalities

The pioneering work of MolT5 has paved the way for research in molecular captioning and generation by description, introducing the ChEBI benchmark dataset for this purpose[18]. Subsequent models such as MoSu[22], MolXPT[23], BioT5[24], and Mol-instruction[25] have expanded the scope of tasks to include numeric molecular property prediction. However, the scarcity of language-molecule pair datasets remains a challenge. To address this, the PubchemSTM[19] and L+M-24[26] datasets have been introduced, leading to improvements in molecular retrieval and editing constrained by language.

### 2.2 Cross Language-Protein Modalities

ProteinDT[27] and Mol-Instruction[25] are examples of multi-modal frameworks that leverage semantically related text for protein annotation and design. BioTranslator[28], a cross-modal model, is specifically designed for annotating biological entities such as gene expression vectors, protein networks, and protein sequences based on user-provided text. Building on the blip2 framework, Mistral and ESM2 have been used to create FAPM[29], which has achieved state-of-the-art results in protein functional Go Terms prediction and demonstrates strong generalization to proteins with few homologs.

### 3 SciMind

In this section, we will detail the design and training of our multi-modal mixture-of-experts model, SciMind. The overview of the pre-training is illustrated in Figure 2. Unlike existing models, our focus is on integrating all modalities into a single model. To this end, we have designed specialized token sets for each modality. However, each modality has a different level of complexity and requires a different number of parameters to avoid overfitting. To leverage the many open-source pretrained language models, we have chosen to construct a Mixture-of-Experts model by splitting the pretrained LLAMA-2-7B model into 16 experts at each of the feedforward layers.

**Figure 2:**
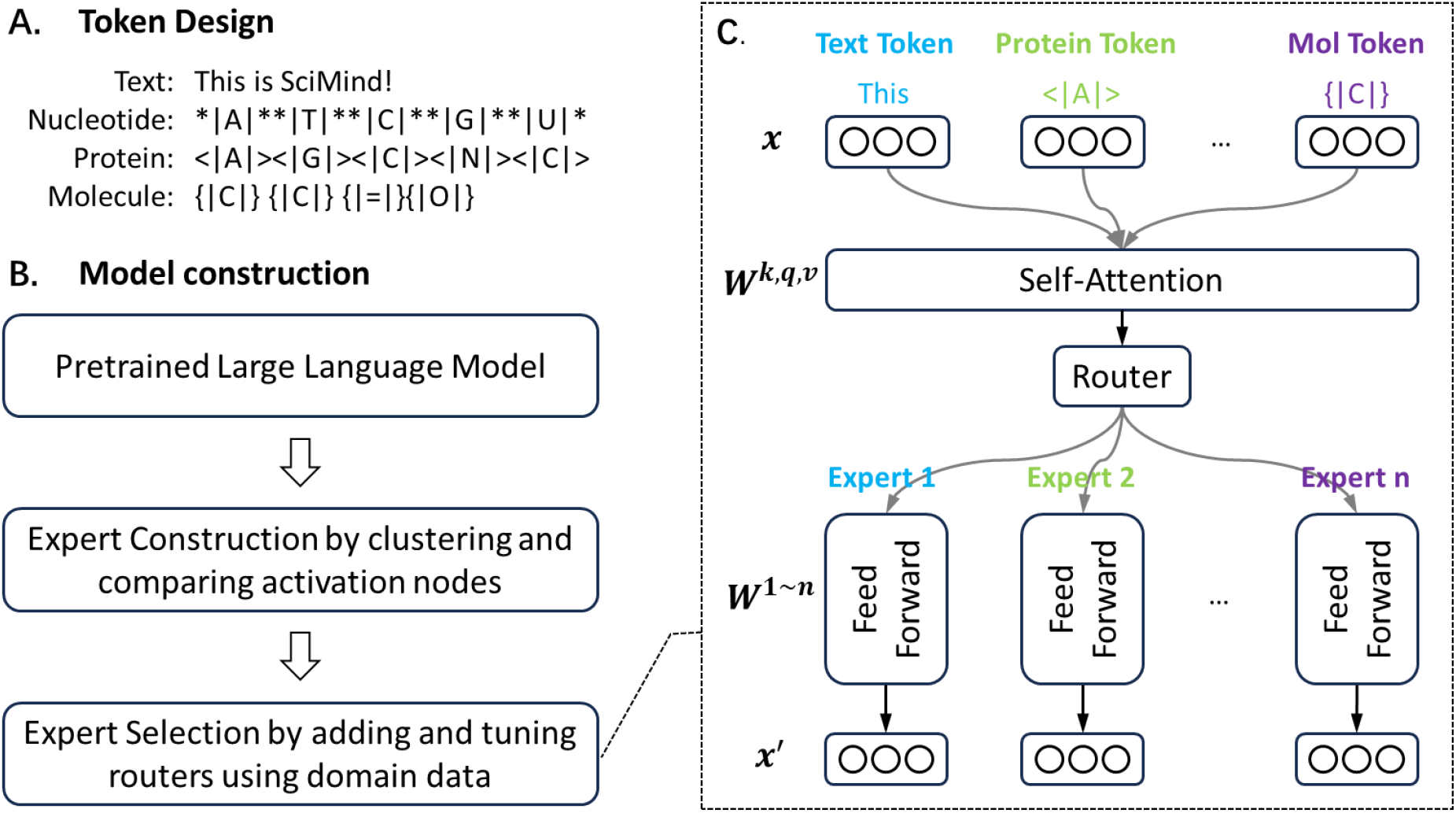
SciMind multi-modal model overview. A, there are four modalities in SciMind, and different modality was designated with different tokens to represent their sequences; B, based on llama-2-7B, 16 experts are split using restricted K-Means clustering according to the feedforward layer weights. A routing layer is added before the feedforward layer of the original model, and domain data is used to pretrain or fine-tune the routing layer to achieve the selection of different experts for different tokens.

### 3.1 Pre-training Corpus

The pre-training corpus includes only single modality data, which are general text, nucleic acids sequences, protein amino acid sequences, and molecule SMILES (Simplified Molecular-Input Line-Entry System) strings. The details of the corpus are provided in Appendix A.

### 3.2 Tokenization

In previous work on cross-language modalities with nucleic acids, molecules, and proteins, the token set was often inherited from NLP methods such as SentencePiece[30]. However, given the different modalities and their unique next-token distributions, we have chosen to tokenize the sequences from nucleic acids, molecules, and proteins by characters, with different brackets used to distinguish characters in different modalities (Figure 2a).

### 3.3 Mixture-of-Experts

Based on the LLAMA-2-7B model, we have split 16 experts using restricted K-Means clustering according to the feedforward layer weights (Figure 2b). A routing layer has been added before the feedforward layer of the original model, and different modality data are fed to pretrain or fine-tune the routing layer to achieve the selection of different experts for different tokens. Considering the propensity to overfit on nucleic acids, protein sequences, and molecule SMILES strings, and our desire to preserve the original language capabilities, we adopted a selective expert activation strategy. For text tokens, we engaged 8 out of the 16 experts. Conversely, for tokens corresponding to other modalities, we restricted the activation to merely 2 out of the 16 experts.

### 3.4 Pretraining

We employed the Huawei MindSpore training framework for pre-training purposes on Huawei Ascend 910 AI chips. Prior to inputting the processed data into the model, an extra step was taken to expedite the training process. This involved converting the data format into the MindRecord format. The Ascend AI framework offers a variety of parallel training modes, efficient memory reuse, and features like automatic mixed precision. These capabilities significantly enhance the training of large-scale models. For further acceleration, we utilized the MindFormer operator during the training process.

## 4 Experiments and Results

### 4.1 Domain Knowledge comprehension

GPT3.5, when utilizing few-shot prompts, tends to struggle with understanding pharmaceutical domain knowledge, particularly in tasks such as name entity recognition and relation extraction. In light of previous research, we evaluated SciMind’s performance on domain knowledge comprehension benchmarks. As illustrated in Table 1, across ten tasks encompassing name entity recognition, relation extraction, and question answering, SciMind surpassed the previous state-of-the-art model, BioLinkBert-Large, in eight tasks.

**Table 1:** Performances on pharmaceutical sciences domain knowledge comprehension and extraction. The metrics of BioLinkBERT-Large and ChatGPT(few-shots) are taken from the original papers.

| Tasks | Entitytype | No.entities | EvaluationMetrics | BioLinkBERT<br>-Large | GPT3.5<br>(few-shots) | SciMind |
| --- | --- | --- | --- | --- | --- | --- |
| **Name Entity Recognition** |  |  |  |  |  |  |
| BC5CDR Disease | Disease | 19,665 | F1entity-level | 0.940 | 0.603 | **0.957** |
| BC5CDR Chem | Chemical | 12,694 | F1entity-level | 0.864 | 0.518 | **0.881** |
| NCBI Disease | Disease | 6881 | F1entity-level | **0.888** | 0.505 | 0.855 |
| BC2GM | Chemical | 79,842 | F1entity-level | 0.852 | 0.375 | **0.898** |
| JNLPBA | Gene | 20,703 | MicroF1 | 0.801 | 0.413 | **0.842** |
| **Relation Extraction** |  |  |  |  |  |  |
| Chemprot | Protein-chemical | 10,031 | MicroF1 | 0.800 | 0.342 | **0.861** |
| DDI | Chemical-chemical | 4,920 | MicroF1 | 0.834 | 0.516 | **0.844** |
| GAD | Gene-disease | 5330 | MicroF1 | **0.849** | 0.524 | 0.805 |
| **Question Answering** |  |  |  |  |  |  |
| PubMedQA | Yes/No/Maybe | 1000 | Accuracy | 0.722 | 0.765 | **0.796** |
| BioASQ | Summary | 885 | Accuracy | 0.948 | 0.886 | **0.950** |

### 4.2 DNA promoter prediction

Predicting gene function is vital for comprehending intricate biological processes. This involves forecasting functional elements and interaction modalities in both coding regions and non-coding sequences that govern gene transcription. Promoters, integral elements in the non-coding regions of genes, regulate gene transcription by managing RNA polymerase binding and initiation. Therefore, the precise prediction of promoter sites is essential for understanding gene expression and genetic regulatory networks.

We evaluated the performance of SciMind using the benchmark data set by DeePromoter. The results presented in Table 2 indicate that SciMind’s predictive performance is on par with DeePromoter in this task. Moreover, SciMind exhibits a slight edge in predicting data with non-TATA promoters. These promoters are more prevalent in certain organisms and types of genes, and they can be involved in more complex regulatory processes.

**Table 2:**
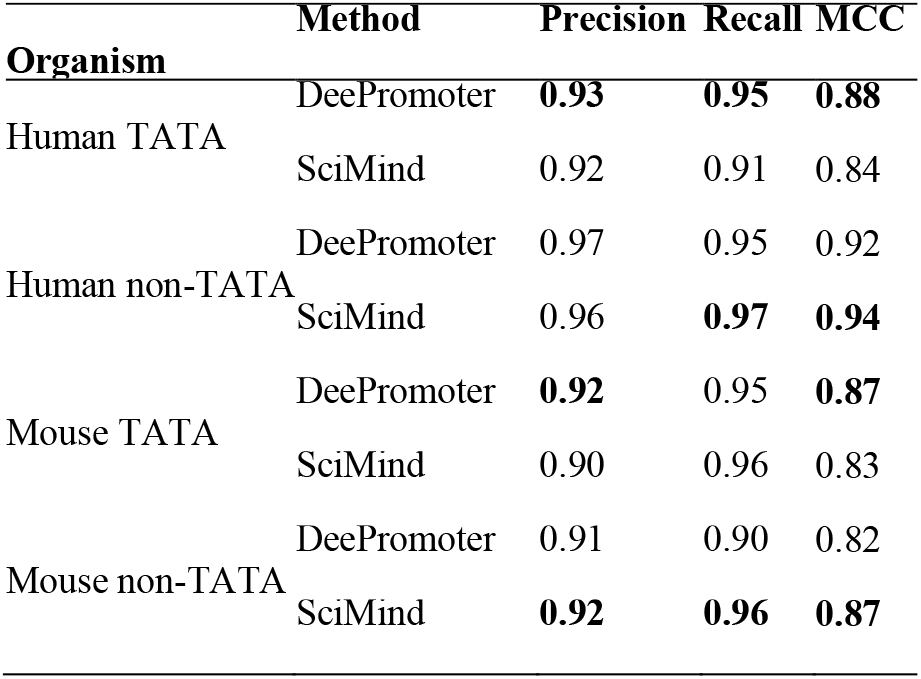
Performances on prompt DNA promoter prediction.

### 4.3 Molecular captioning

The objective of the molecule captioning task is to provide a structural or biological functional description for a given molecule. In our approach, we represent molecules using SMILES strings, thereby transforming the task into a seq2seq translation problem. This problem is well-suited for processing by large language models. We have two benchmark datasets with varying sizes. The ChEBI dataset is annotated by humans, while the L+M-24 dataset is summarized by ChatGPT. A notable difference is that some ChEBI data includes descriptions identifying the core structures of molecules.

As shown in Table 3, our Mixture-of-Experts-based SciMind model achieves state-of-the-art (SOTA) performance on most of the metrics in both benchmark datasets.

**Table 3:**
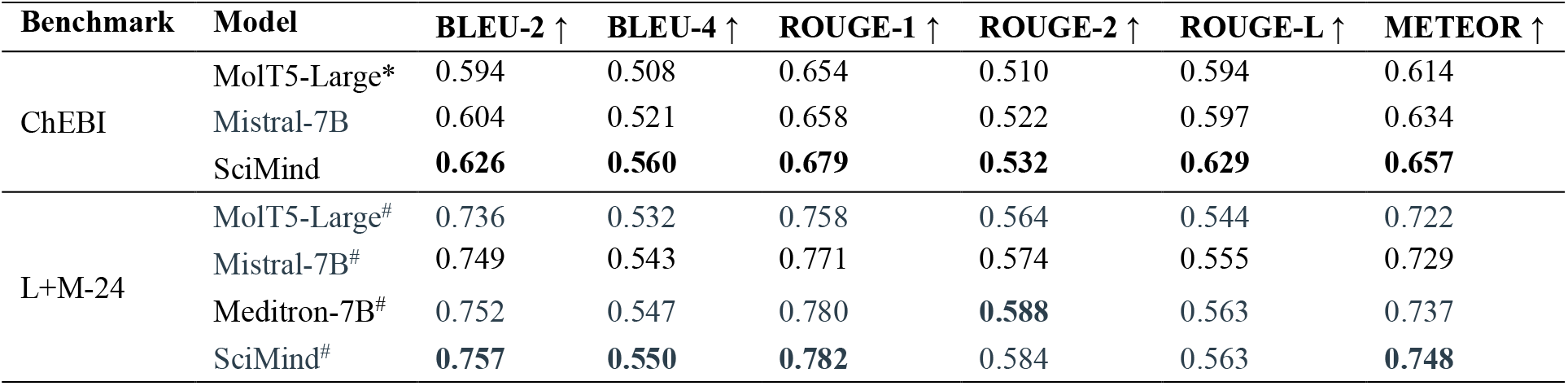
Performances on molecular captioning. The metrics value of methods annotated with * are taken from the original paper. And the metrics value of methods annotated with ^#^ are taken from the contest leaderboard (https://www.codabench.org/competitions/2914), where SciMind ranked No.1. Other metrics values are evaluated following the process of previous work.

| Benchmark | Model | BLEU-2 ↑ | BLEU-4 ↑ | ROUGE-1 ↑ | ROUGE-2 ↑ | ROUGE-L ↑ | METEOR ↑ |
| --- | --- | --- | --- | --- | --- | --- | --- |
| ChEBI | MolT5-Large* | 0.594 | 0.508 | 0.654 | 0.510 | 0.594 | 0.614 |
|  | Mistral-7B | 0.604 | 0.521 | 0.658 | 0.522 | 0.597 | 0.634 |
|  | SciMind | **0.626** | **0.560** | **0.679** | **0.532** | **0.629** | **0.657** |
| L+M-24 | MolT5-Large<sup>#</sup> | 0.736 | 0.532 | 0.758 | 0.564 | 0.544 | 0.722 |
|  | Mistral-7B<sup>#</sup> | 0.749 | 0.543 | 0.771 | 0.574 | 0.555 | 0.729 |
|  | Meditron-7B<sup>#</sup> | 0.752 | 0.547 | 0.780 | **0.588** | 0.563 | 0.737 |
|  | SciMind<sup>#</sup> | **0.757** | **0.550** | **0.782** | 0.584 | 0.563 | **0.748** |

### 4.4 Molecular generation

Molecular generation is the reverse task of molecule captioning. Given a natural language description of the desired molecule, the goal is to generate a molecule that matches the description. The results in Table 4 demonstrate that our Mixture-of-Experts-based SciMind achieves state-SOTA performance on most metrics across both benchmarks.

**Table 4:**
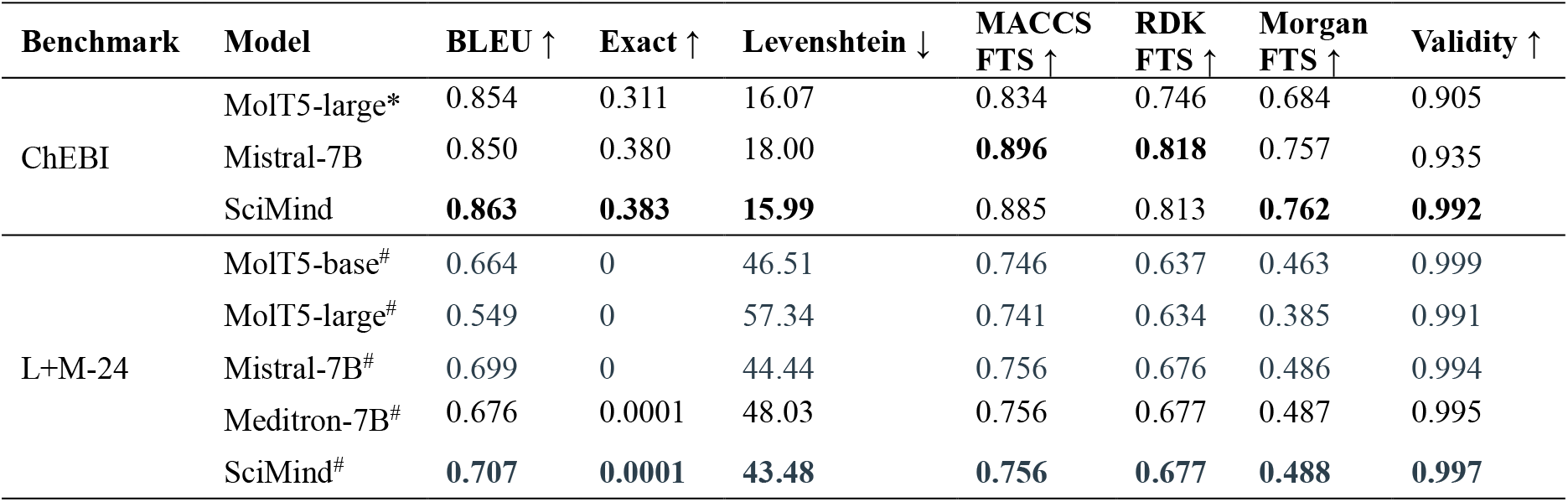
Molecular generation based on description. The metrics value of methods annotated with * are taken from the original paper. And the metrics value of methods annotated with ^#^ are taken from the contest leaderboard (https://www.codabench.org/competitions/3014), where SciMind ranked No.1. Other metrics values are evaluated following the process of previous work.

### 4.5 Protein-oriented prediction

We leverage the protein-oriented instruction dataset from Mol-Instruction to fine-tune SciMind. Figure 3 shows the Rouge-L metrics of five methods across four tasks: protein function, general description, catalytic activity, and domain/motif prediction. Compared to other single-modal language models, SciMind achieves the best performance on Rouge-L metrics across all four tasks.

**Figure 3:**
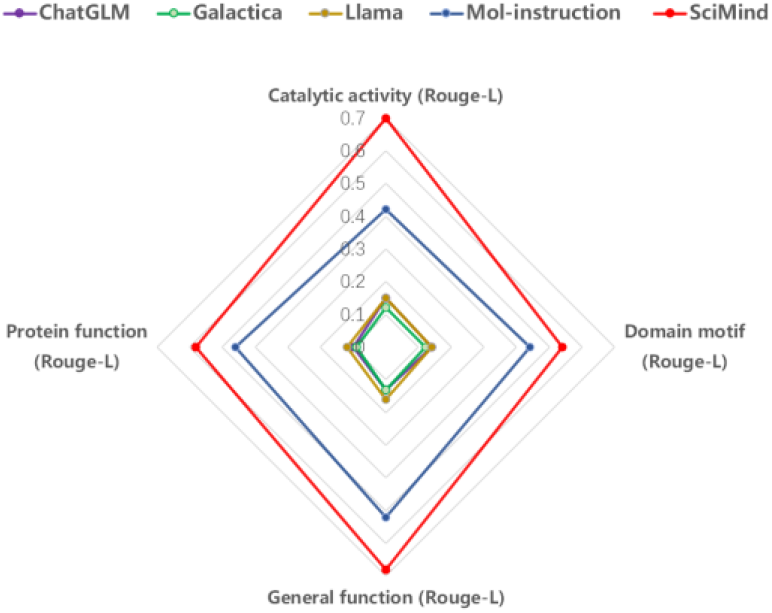
Performance on protein-oriented prediction

## 5 Conclusions and discussion

In this paper, we introduced SciMind, a unified pre-training framework designed to include all the modalities in pharmaceutical sciences. we’ve designed a specialized token set and introduce a new pre-training and fine-tuning strategy that leverages the advantages of large-parameter models while minimizing their expenses. This strategy, supported by a prior expert allocation and selection mechanism, allows data of different modalities to choose the most suitable processing path. This method not only leads to a sparser model architecture, thus cutting down on inference costs, but also avoids modal alignment and the potential performance decrease due to model size reduction. We’ve created a multi-modal mixture-of-expert foundational large model for pharmaceutical sciences, named SciMind. This model has undergone extensive pre-training on publicly accessible datasets including nucleic acids, protein sequences, molecular structures strings, and biomedical texts, and could be fine-tuned for downstream tasks involving all modalities in pharmaceutical sciences. The experimental outcomes suggest that the SciMind model not only delivers outstanding performance but also shows high flexibility and interpretability in response to prompt words, offering a sturdy base for its use in pharmaceutical sciences.

Due to the lack of well-aligned multimodal data, our model has not fully demonstrated its advantages. In addition to molecular captioning and generation by description, the inclusion of the protein modality will make the interaction between the language and small molecule modalities more explainable and useful. This approach helps accumulate more information and is a promising direction to explore.

## Acknowledgments

We acknowledge the support by the Huawei MindSpore team.

## A Pre-training Corpus

### DNA data

Our pretraining data for nucleic acid sequences is derived from DNABERT_S, which includes a human genome dataset containing 2.75 billion nucleotide bases. The multi-species genome dataset includes genomes from 135 different species, distributed across 6 categories and containing a total of 32.49 billion nucleotide bases, which is 12 times the size of the human genome dataset. We use *| |* to separate the characters in the nucleic acids, as shown in Figure 2a.

### RNA data

This dataset is a subset of the RNAcentral active fasta file, available at <https://ftp.ebi.ac.uk/pub/databases/RNAcentral/releases/24.0/sequences/rnacentral_active.fasta.gz>, that has been converted to the parquet format. It represents approximately 10% of the overall dataset and contains 3,252,483 (3.2 million) sequences, comprising a total of 2,642,703,990 (2.6 billion) bases. We use *| |* to separate the characters in the nucleic acids, as shown in Figure 2a.

### Protein data

Protein sequence databases, such as UniParc, contain a wide variety of sequences from different organisms. In our experiments, we follow the esm work and used the 250 million sequences from the UniParc database, which contains a total of 86 billion amino acids. These datasets are similar in size to large text corpora that are commonly used to train high-capacity neural network models for natural language processing tasks. We use <| |> to separate the characters in the protein sequences, as shown in Figure 2a.

### Molecule data

The molecular data is taken from https://huggingface.co/datasets/kjappelbaum/chemnlp_iupac_smiles, which contains 30 million molecules’ SMILES and their IUPAC names. We use {| |} to separate the characters in the molecule SMILES, as shown in Figure 2a.

## B Finetuning corpus

All downstream tasks in this paper have been benchmarked against previous studies. Accordingly, we fine-tune and test our models using either the pre-split datasets or by splitting the data in the same manner as the original studies.

